# Major urinary protein (*Mup*) gene family deletion drives sex-specific alterations on the house mouse gut microbiota

**DOI:** 10.1101/2023.08.01.551491

**Authors:** Madalena V. F. Real, Melanie S. Colvin, Michael J. Sheehan, Andrew H. Moeller

## Abstract

The gut microbiota is shaped by host metabolism. In house mice (*Mus musculus*), major urinary protein (MUP) pheromone production represents a considerable energy investment, particularly in sexually mature males. Deletion of the *Mup* gene family shifts mouse metabolism towards an anabolic state, marked by lipogenesis, lipid accumulation, and body mass increases. Given the metabolic implications of MUPs, they may also influence the gut microbiota. Here, we investigated the effect of deletion of the *Mup* gene family on the gut microbiota of sexually mature mice. Shotgun metagenomics revealed distinct taxonomic and functional profiles between wildtype and knockout males, but not females. Deletion of the *Mup* gene cluster significantly reduced diversity in microbial families and functions in male mice. Additionally, specific taxa of the Ruminococcaceae family, which is associated with gut health and reduced risk of developing metabolic syndrome, and several microbial functions, such as transporters involved in vitamin B5 acquisition, were significantly depleted in the microbiota of *Mup*-knockout males. Altogether these results show that major urinary proteins significantly affect the gut microbiota of house mouse in a sex-specific manner.

**Importance:** The community of microorganisms that inhabit the gastrointestinal track of animals, known as the gut microbiota, can have profound effects on host phenotypes. The gut microbiota is in turn shaped by host genes, including those involved with host metabolism. In adult male house mice, expression of the major urinary protein (*Mup*) gene cluster represents a substantial energy investment, and deletion of *Mup* gene family leads to fat accumulation and weight gain in males. We show for the first time that deleting *Mup* genes also alters the gut microbiota of male, but not female, mice in terms of both taxonomic and functional composition. Male mice without *Mup* genes harbored fewer gut bacterial families and reduced abundances of several species, including bacteria previously shown to reduce obesity risk. Studying the impact of the *Mup* genes on the gut microbiota will help us understand how these genes influence host phenotype more broadly.

## Introduction

The gut microbiome has emerged as a major modulator of host phenotypes, from metabolism [1-5] to behavior [6-8], motivating the investigation of the factors that shape the gut microbial community. Hosts can influence the taxonomic and functional composition of the microbiota through various genetically based physiological processes [9]. Gene knockouts allow us to test hypotheses concerning the effects of these processes on the gut microbial community. This approach has been employed to demonstrate the effects on the microbiota of innate and adaptive immune genes [10-14]. However, the effects on the microbiota of non-immune genes related to host metabolism have only recently started to be investigated [15-17].

Major urinary proteins (MUPs) are lipocalins involved in pheromonal communication [18-21], and their production represents a major metabolic investment for house mice (*Mus musculus*). In this social rodent, the *Mup* gene family has undergone an extensive parallel evolutionary expansion, accumulating 21 distinct copies in a 2.2 Mbp gene cluster on chromosome 4 [21]. The genetic diversity and dynamic expression of *Mup* genes allow excreted MUPs to function as individual identifiers [22-25], conveying kinship, territory, social status, sex, reproductive state, age, health, and even diet [26-31]. This communication occurs mostly through urine markings [32, 33], with MUPs constituting up to 90% of the male urinary proteome [34], a 2–8 times higher protein content than females [35]. *Mup* genes are also the most highly expressed genes in the liver [35], representing up to 20% of the hepatic transcriptome in mature males [36]. Male MUP expression is particularly upregulated after puberty [37, 38], when social dominance is established [39, 40].

MUP production is a considerable energy investment for mice, particularly males. MUP expression is reduced under caloric restriction [41-43] and in obese and diabetic mice [41, 44]. In addition to being affected by energy availability, MUPs also regulate house-mouse metabolism. Genetically obese and diabetic mice inoculated with a recombinant MUP display improved insulin sensitivity mediated by a reduction in glucose and lipid anabolism [44]. Conversely, sexually mature *Mup*-knockout (KO) males exhibit increased anabolic phenotypes relative to wildtype (WT) individuals [45]. KO males displayed higher body weight and visceral adipose tissue than WT mice, despite lower food intake and equal energy expenditure. This metabolic shift also manifested through higher circulating levels of triglycerides, free fatty acids and leptin, and an upregulation of genes associated with lipid metabolism in KO *versus* WT males. These results point to the profound effect of MUPs on mouse metabolism. Given the known interactions between host metabolic function and the gut microbiota [11, 15-17, 46, 47], *Mup* expression may indirectly impact the gut microbial community. Additionally, *Mup* gene expression has been observed in the intestinal transcriptome of juvenile males [48] and in the duodenal proteome of adults [49], indicating gut commensals could also be in direct contact with MUPs. These possible mechanisms lead us to hypothesize that deletion of the *Mup* gene family may have major effects on the house-mouse microbial community, but this has yet to be tested.

Here, we investigated how deletion of the *Mup* gene cluster impacts the gut microbial community of house mice. To answer this question, we generated a *Mup*-knockout line (KO; *Mup*^-/-^) with CRISPR/Cas9 and crossed it with wildtype mice (WT; *Mup*^+/+^) for multiple generations, yielding litters of mice discordant for *Mup* genotype. We then sequenced and compared the gut metagenomes of the homozygous progeny (KO vs WT). We hypothesized that sexually mature KO and WT mice would host distinct microbiotas, both taxonomically and functionally, and that the largest differences would be found in males. We found a sex-specific effect of the *Mup* knockout on the microbial taxonomic and functional profiles, demonstrating this metabolically costly gene shapes the gut microbiota of house mice.

## Results

### Deletion of the Mup gene cluster

The *Mup* gene cluster was fully deleted using CRISPR/Cas9 to cleave upstream of *Mup*4 and downstream of *Mup*21 (Figure 1A). An individual’s *Mup* status was confirmed by genotyping ear biopsies collected at weaning. Lack of MUP production was confirmed by measuring urinary MUP levels (Figure 1B). Heterozygous crosses generated offspring with two copies of the *Mup* gene cluster (wildtypes, WT), only one (heterozygotes, HT), or none (*Mup* knockouts, KO) at the expected Mendelian ratio (Figure S1).

**Figure 1.**
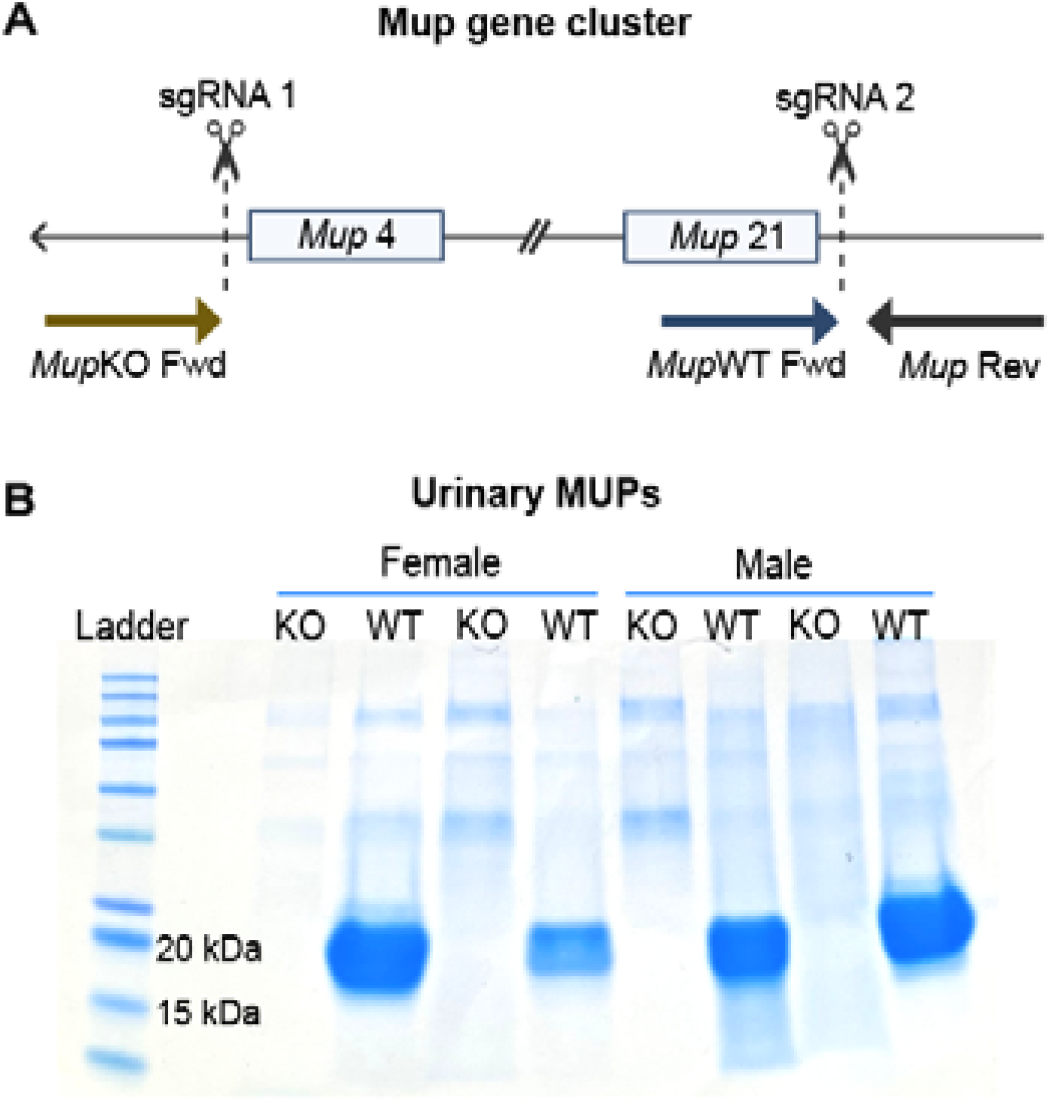
*Mup* deletion and experiment timeline. (A) Representation of the 2 Mbp *Mup* gene cluster shows the CRISPR/Cas9 sgRNA target sites (dashed lines) and the primer binding sites used for genotyping (arrows). (B) Representative SDS-PAGE stained with Coomassie Brilliant Blue shows urinary MUPs in WT mice (blue band at 20 kDa) and the absence of MUPs in KO individuals of both sexes (n = 2 mice/sex/genotype).

### Metagenomic sequencing of Mup KO and WT mice

MUP production is upregulated in WT males after sexual maturity [37, 38] and thus we sampled the gut microbiota by collecting fecal pellets from 12-week–old mice (87 ± 3 days old) housed in holding cages with same-sex littermates of diverse genotypes. A total of six litters were sampled. We sequenced the fecal metagenome of WT and KO mice, yielding an average of 33.6 ± 20.1 million reads/sample and 33.4 ± 20.1 million reads/sample post-quality control and host-filtering. For the alpha and beta diversity analyses, all samples were rarefied to the minimum observed value of 1 million reads/sample. Functional annotation of unrarefied reads produced 478.1 ± 295.9 thousand COG counts/sample, which were rarefied to the minimum observed value of 84.0 thousand COG counts/sample before alpha and beta diversity analyses.

### Mup genotype affected taxonomic composition of the gut microbiota of sexually mature males

To test the hypotheses that *Mup* genotype affected the composition of the gut microbiota, particularly in males, we compared the gut microbiota of sexually mature KO and WT mice. PERMANOVA analyses based on Jaccard and Bray-Curtis dissimilarities among samples’ MetaPhlAn 4 taxonomic profiles revealed that the microbiota of males and females was significantly different (Table S1). Thus, we tested for significant differences between KO and WT individuals within each sex while controlling for litter effects. These analyses revealed that mature WT and KO males display significantly different microbial taxonomic compositions at the species and genus (but not family) levels based on both presence-absence (Jaccard) and abundance-weighted (Bray-Curtis) beta diversity dissimilarities, even while co-housed with same-sex littermates of diverse *Mup* genotypes (Table 1; Table S2; Figure S2). This significant shift in the taxonomic makeup of the microbiota between WT and KO males was not driven by differences in dispersion between genotypes, as no significant differences in taxonomic PERMDISP were observed (Table 1; Figure 2B). No significant differences were observed in females (Table 1; Table S2; Figure S2). Cumulatively, these results show that deletion of the *Mup* gene caused a sex-specific directional shift in the taxonomic composition of the gut microbiota.

**Figure 2.**
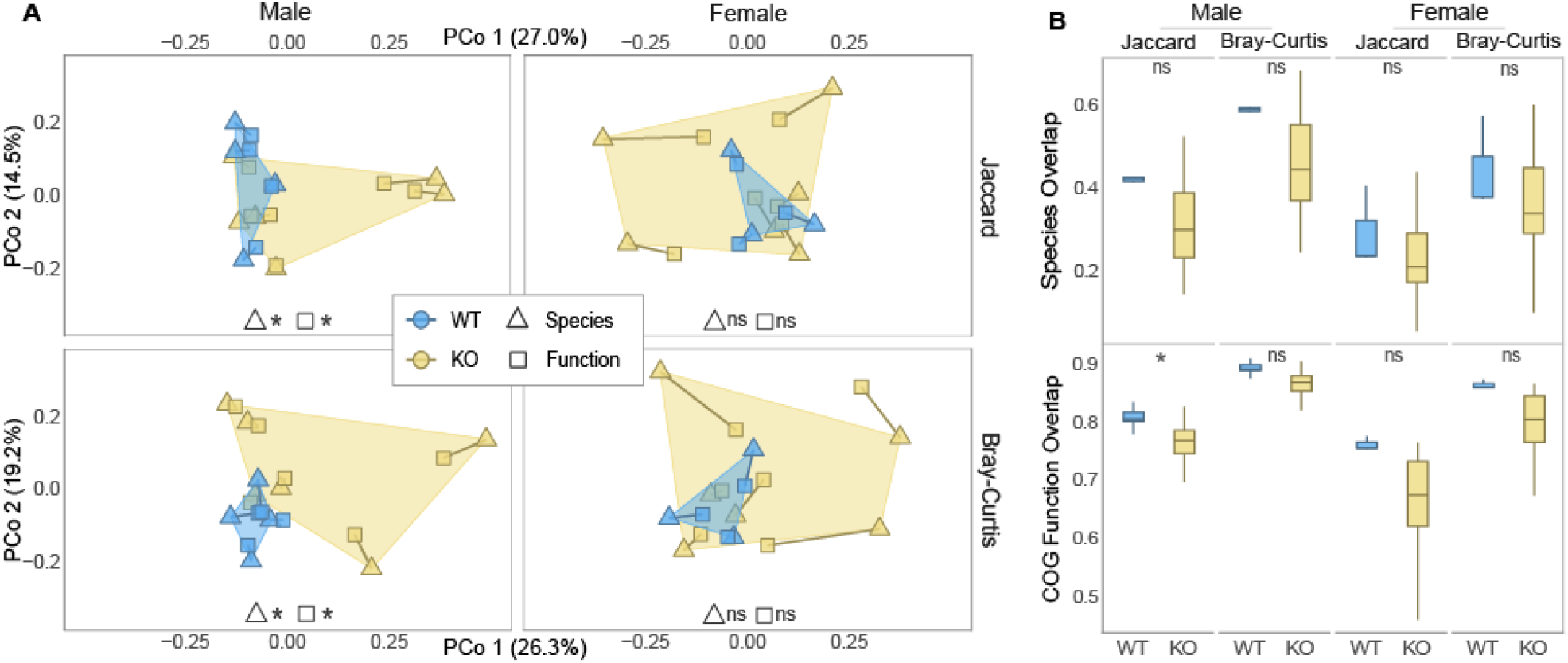
*Mup* deletion significantly changes the gut microbial taxonomic and functional composition of mature males. (A) Principal coordinates analyses (PCoA) show the ordinated Jaccard (top row) and Bray-Curtis (bottom row) dissimilarities in species (triangles) and COG function (squares) composition in the fecal metagenome of sexually mature mice, superimposed with a Procrustes analysis. The ordinated points are faceted by sex and colored by genotype, showing the taxonomic and functional profiles of WT (blue) and KO (yellow) male (left column) and female (right column) mice. The percentage of variation in the taxonomic dissimilarity matrix explained by each PCo axis is enclosed in parenthesis. The PERMANOVA analyses (center bottom) indicate whether there is a significant difference in the centroid and/or dispersion of the WT and KO groups for each dissimilarity measure and sex (* = *p*-value < 0.05; ns = *p*-value > 0.05). (B) Box plots show the species (top row) and COG function (bottom row) overlap in the microbiota of mature males (left column) and females (right column), using both Jaccard (left sub-column) and Bray-Curtis (right sub-column) similarities. The PERMDISP analyses (center top) indicate whether there is a significant difference in the dispersion of the WT (blue) and KO (yellow) groups (* = *p*-value < 0.05; ns = *p*-value > 0.05).

**Table 1.** Effect of *Mup* genotype on microbial species and COG function composition^a^.

| *Sex* | *Dissimilarity* | *Species* |  | *COG Function* |  |
| --- | --- | --- | --- | --- | --- |
|  |  | *PERMANOVA* | *PERMDISP* | *PERMANOVA* | *PERMDISP* |
| *Male* | *Jaccard* | **0.037** | 0.273 | **0.021** | **0.046** |
|  | *Bray-Curtis* | **0.019** | 0.188 | **0.012** | 0.080 |
| *Female* | *Jaccard* | 0.154 | 0.739 | 0.116 | 0.633 |
|  | *Bray-Curtis* | 0.158 | 0.398 | 0.481 | 0.200 |

<sup>a</sup> Significant results are **bolded** (Holm–Bonferroni adjusted *p*-value < 0.05).

### Mup genotype affected functional composition of the gut microbiota of sexually mature males

We also tested if gene function profiles in the microbiota differed between mouse genotypes. Using MG-RAST annotations based on Clusters of Orthologous Genes (COGs) (Table S2), we conducted the same PERMANOVA and PERMDISP analyses that were employed for the microbiota taxonomic profiles. As observed for taxonomy, the COG functional composition was significantly different between WT and KO males, but not between females (Table 1; Figure S3). In contrast with the taxonomic results, PERMDISP indicated higher functional variation among KO males compared to WT individuals, based on Jaccard, but not Bray-Curtis, similarities (Table 1; Figure 2B). No such differences in PERMDISP were observed in females (Table 1; Figure 2B). These findings indicate mature KO males exhibited both directional shifts and increased inter-individual heterogeneity in the functional composition of the gut microbiota relative to WT mice.

### Significant correspondence of taxonomic and functional profiles

Based on the results of PERMANOVA and PERMDISP analyses (Table 1), we next visualized the similarities among the microbiota of WT and KO mice at both taxonomic and functional levels using Procrustes. These analyses revealed significant correspondence between taxonomic and functional profiles recovered from individual mice using Bray-Curtis but not Jaccard (Table S3). Within sexes, only in mature males did the taxonomic and functional configurations significantly matched (Table S3). Plotting the superimposed taxonomic and functional compositions of mature mice revealed that samples clustered by *Mup* genotype in males but not females (Figure 2A).

### Gut microbial alpha diversity varied between Mup genotypes

Given the observed differences in gut taxonomical and functional profiles between WT and KO males, we investigated whether the *Mup* knockout also affected taxonomic and functional alpha diversity. A linear mixed-effects model accounting for litter as a random effect indicated that KO males had lower diversity (Shannon) and evenness (Pielou) of microbial families, but not species or genera, than their WT counterparts (Table S4; Figure S4). Females displayed no effect of *Mup* genotype on taxonomic diversity (Table S4; Figure S4). We also observed differences in COG function diversity (Shannon, but not Pielou) between WT and KO males (Table S4; Figure S5). No such differences were observed in females (Table S4; Figure S5). These alpha diversity results align with our findings from the beta diversity analyses, once again showing that the *Mup* gene cluster deletion affected the gut microbiota diversity of mature mice in a sex-specific manner.

### Specific microbial taxa and functions differed in abundance between Mup genotypes

Given that the *Mup* knockout affected the gut microbial taxonomic and functional composition in mature males (Figure 2), we next identified which specific taxa and/or COG functions differed between genotypes (Figure 3 and S6). Differential abundance analyses with ANCOM-BC2 detected a Ruminococcaceae species, GGB30296 SGB43260, that was significantly underrepresented in mature KO males (Holm–Bonferroni adjusted *p*-value < 0.001; Table S5). No taxa were differentially abundant at this significance threshold in females (Table S5). Several COG functions were underrepresented in KO males (Table S5). The largest fold-change in abundance was observed in the Na+/panthothenate symporter, involved in the transport of panthothenate or vitamin B_5_ [50]. The most significant was component EscU of the Type III secretory pathway, used by Gram-negative bacteria to inject virulence factors into host cells [51, 52]. L−asparaginase II, a periplasmic high-affinity enzyme that hydrolyses exogenous l-asparagine into l-aspartate and ammonia [53], was also highly significant. A hypergeometric test revealed that no single COG category was particularly overrepresented among the various depleted functions in KO males (Table S6). The microbiota of KO males was instead significantly enriched in transcriptional regulators. Here too, no functions were highly differentially abundant in the microbiota of females (Table S5). These results point to the sex-specific effects of the *Mup* gene cluster deletion, not just at the level of the whole microbial community but affecting the abundance of specific taxa and functions too.

**Figure 3.**
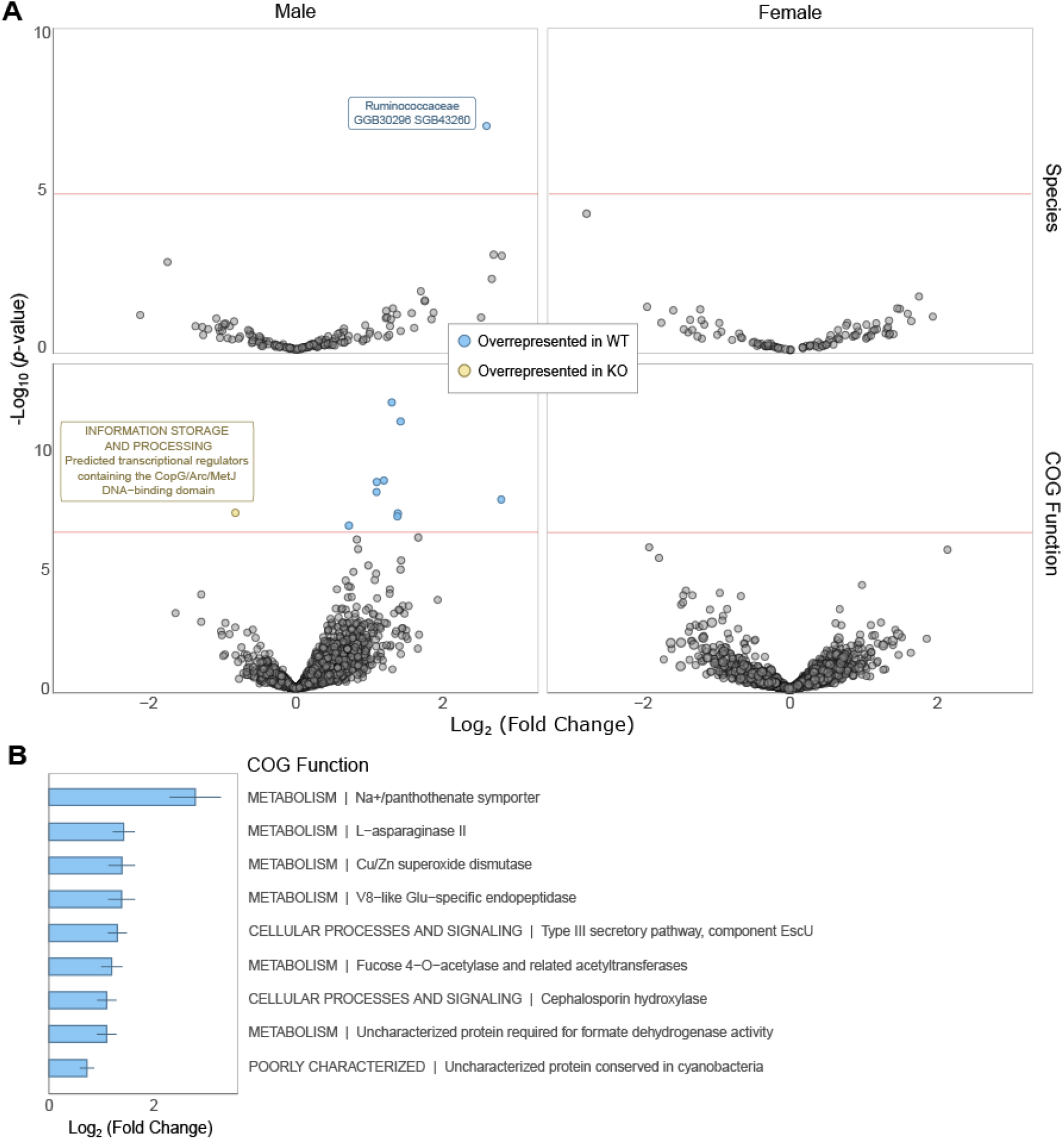
*Mup* deletion significantly shifts the abundance of various microbial taxa and functions. (A) Volcano plots show the log_2_-transformed fold change (LFC) in the abundance of species (top row) and COG functions (bottom row) in the gut microbiota of mature male (left column) and female (right column) mice. ANCOM-BC2 analyses identified species and functions (points) that were significantly more abundant in WT (blue) or KO mice (yellow), labeled respectively with the family and species designation, and the COG Category and Function. Red lines mark the significance threshold (Holm–Bonferroni–adjusted *p*-value < 0.001). The y-axis indicates the -log_10_ transformation of the non-adjusted *p*-value. (B) Bar plots show the LFC in abundance of the COG functions that were significantly overrepresented in mature WT males.

## Discussion

We found that the presence of the *Mup* gene cluster in house mice significantly altered the taxonomic and functional composition of the gut microbial community. In accordance with our hypothesis, deletion of the *Mup* gene cluster significantly affected the gut microbiota of mature male mice, but not female mice, through a shift in composition, reduction in diversity, and depletion of microbial taxa and functions. These differences were seen even among co-housed littermates, indicating that the effects of *Mup* were robust to the homogenizing effect of microbial dispersal among animals sharing a cage [54, 55]. Furthermore, *Mup* deletion did not change the microbial profiles of females. This aligns with our hypothesis that the effect of a *Mup* deletion would be stronger in male mice, given the sexually dimorphic expression pattern this gene cluster exhibits [35, 36].

The *Mup* deletion led not only to a shift in the taxonomic and functional composition of mature males, but WT individuals tended to display higher inter-individual homogeneity. This pattern reached significance at the functional level. The production and/or presence of MUPs at high levels could exert a parallel pressure on the microbial community of WT males, making it converge on a similar configuration. Mature males lacking the *Mup* gene cluster also exhibited decreased diversity and evenness in microbial families and gene functions. A reduction in microbial taxonomic and functional diversity is one of the hallmarks of obesity-related metabolic syndrome [3, 56, 57], and *Mup* KO males were previously shown to develop phenotypes associated with this syndrome, such as higher body weight, visceral adipose tissue, and circulating levels of triglycerides, free fatty acids and leptin [45]. Cumulatively, these results are consistent with a scenario in which knocking out the *Mup* gene cluster dysregulates mouse development and/or physiology, increasing the amount of variation in functional content among individuals and driving reductions in alpha diversity.

In addition to community-level patterns, the abundance of specific taxa was impacted by the deletion of the *Mup* gene cluster. Particularly, a Ruminococcaceae species was significantly depleted in KO males. This microbial family has been previously associated in both mice and humans with lower body weight and reduced risk of developing metabolic syndrome [58-61]. Ruminococcaceae are major butyrate producers [62], a short-chain fatty acid that promotes gut health and increases satiety [63]. The depletion of this taxa in *Mup* KO males might be associated with the previously observed weight gain in these animals [45], although further experiments (e.g., microbiota-transplant experiments into germ-free mice) will be required to assess this hypothesis.

Several gene functions were also depleted in the gut microbiota of *Mup* KO males, particularly the sodium-dependent transporter of panthothenate (vitamin B_5_) [50]. Vitamin B_5_ forms the core of coenzyme A (CoA), an essential co-factor in cellular respiration [64] and energy metabolism, such as the synthesis of fatty acids and complex lipids [65]. Animals need to acquire vitamin B_5_ from their diet or gut commensals [65], but in the gut microbiota only Bacteroidetes and Proteobacteria have been found to synthesize vitamin B_5_ *de novo* [66]. Microbes from non-producing phyla rely on sodium-dependent transporters to acquire this essential vitamin, forming codependence networks with producers [66]. The depletion of the Na^+^/panthothenate symporter in the microbiota of KO males could represent not only a disturbance to these microbial networks but an eventual deficit in vitamin B_5_ for the host.

One potential caveat to the observed sex-dependent effects of *Mup* deletion on the house mouse microbiota is that our study may have been underpowered to detect small effects in females, even if they occurred in our study. However, subsampling the WT males by randomly removing one individual and reperforming all analyses indicated that significant results were still observed in iterations based on lower sample sizes (see Supplemental Results), suggesting that sampling effort alone cannot explain the disparity in results between males and females. Regardless, a priority for future work will be to test through expanded sampling whether WT and KO females also differ in the taxonomic and functional composition of their microbiota.

The observed shifts in the microbial community – through a decrease in overall diversity and depletion of taxa and functions associated with host metabolic health – can in turn profoundly impact the metabolism of KO mice. It will be interesting to disentangle which aspects of the anabolic phenotype observed in *Mup*-knockout males are directly caused by the absence of MUP production or being mediated by the taxonomic and functional shifts in the microbial community. Reciprocally transplanting the gut microbiota of WT and KO mice to germ-free mice with different *Mup* genotypes could elucidate which metabolic phenotypes are sufficiently caused by the microbial community. The role of the identified Ruminococcaceae species in particular could be explored by inoculating KO mice with this taxon and observing changes in the mouse metabolic state. Additionally, it is worth investigating the mechanisms through which deletion of the *Mup* gene cluster affects the gut microbiota. Inoculating KO mice with recombinant MUPs [44] could help differentiate between the effects of MUP production and the role of circulating MUPs on the house-mouse gut microbial community. Fully understanding the interactions between MUPs, the microbiota, and metabolism will reveal the role of this sexually dimorphic gene on house-mouse physiology.

## Methods

All procedures conformed to guidelines established by the U.S. National Institutes of Health and have been approved by the Cornell University Institutional Animal Care and Use Committee (protocol #2015-0060).

### Genome Editing

The *Mup* gene cluster knockout with CRISPR/Cas9 on FVB x B6 hybrid mice was performed by Cornell’s Stem Cell and Transgenic Core Facility. The inbred mouse strain FVB/NJ (JAX #001800) is commonly used to generate transgenics, due to its large pronucleus and litter size [67]. B6(Cg)-*Tyr^c-2J^*/J (JAX #000058) are C57BL/6J albino mice [68]. Purified RNA (Cas9 + sgRNA; sgRNA1: GGGCCATAAGGAATGATCTTGGG; sgRNA2: GAGCTAAAGGAGACCCATATGGG) was injected into the pronucleus and cytoplasm of fertilized FVB x B6 embryos (n = 150). Embryos that advanced to 2-cell stage were transferred into pseudo-pregnant FVB x B6 females (20 embryos per recipient).

The resulting offspring were genotyped with Transnetyx (Cordova, TN) using ear tissue samples collected at weaning. Real-time PCR was used to detect *Mup* presence, using primers that targeted the gene cluster (forward primer: ACAACCTGCCATTCTGTCTCTTAAT; reverse primer: GGCAATGAAACAAGGATTTGAGTTTTACATAT; final concentration: 900 nM). A second test confirmed *Mup* deletion by using primers flanking the gene cluster, as amplification is only possible if the 2.2 mbp region is absent (forward primer: CAGTACTCAGGGCTTGGGATT; reverse primer: ACTGTTCTCGTGGGAATATGTATTGTGAA; final concentration: 900 nM). Successful amplification in both tests indicates a heterozygous (HT) genotype, with *Mup* present in one of the chromosomes and missing in the other. Wildtype (WT) individuals will only have amplification in the detection test, with only the deletion test being successful in knockouts (KO). The genotyping was phenotypically confirmed by measuring MUP concentration in the animals’ urine with sodium dodecyl sulfate–polyacrylamide gel electrophoresis (SDS-PAGE). The gel was stained with Bio-Rad’s Bio-Safe™ Coomassie brilliant blue and we used Bio-Rad’s Precision Plus Protein™ Kaleidoscope™ Prestained Protein Standards as a ladder.

### Animals

A breeding population was kept at a conventional mouse facility at Cornell University, Ithaca, NY. Heterozygous crosses generated litters of WT, HT, and KO individuals. We analyzed mice from 6 different litters (7 ± 1 pups per litter). Mice were weaned at 3 weeks of age (24 ± 3 days) and housed with same-sex siblings of diverse genotypes (2-4 animals in a cage). Animals were kept in a 12h light:12h dark cycle, with constant room temperature and humidity (21°C, 50%). Standard chow diet and water were available *ad libitum*. Only WT and KO males and females were included in the analysis (n = 20; 4 WT males; 3 WT females; 6 KO males; 6 KO females).

### Microbiota analysis

Fecal samples were collected at 12 weeks of age (87 ± 3 days). Microbial total DNA was extracted using the Quick-DNA MagBead extraction kit (Zymo, Irvine, CA) and the OT-2 liquid handling robot (Opentrons, New York, NY). Library preparation followed the Hackflex protocol [69] using the same robot. Libraries were sequenced on an Illumina NextSeq 2000 (Cornell’s Biotechnology Resource Center, Ithaca, NY).

The metagenomic data was quality controlled with FastQC [70], followed by trimming of the Illumina adapters with Cutadapt [71], and removal of host reads with Bowtie2 [72]. The remaining reads were taxonomically profiled with MetaPhlAn4 [73] and functionally annotated with Clusters of Orthologous Croups (COGs) using MG-RAST [74].

All data analyses were conducted in R (v4.2.2). Before measuring alpha and beta diversity, the library size was normalized by randomly subsampling sequences to 1 million reads/sample for the taxonomic data, and 84 thousand COG counts/sample for the functional data. The Shannon diversity index, observed richness and Pielou’s evenness metrics were measured using the microbiome R package (v1.20.0) [75]. The Jaccard and Bray-Curtis dissimilarities and Euclidean distances between samples were calculated with phyloseq (v1.42.0) [76]. All plots were created with ggplot2 (v3.4.1) [77]. All figure panels were assembled using Inkscape 1.2.

### Statistical analysis

Data are presented as mean ± standard deviation. Group means were compared with the Wilcoxon signed-rank test. A linear mixed-effects model assessed the effect of genotype on microbiota diversity, while controlling for litter as a random effect (via lmerTest v3.1 [78]). The factors (genotype, sex and/or litter) that explained the microbiota dissimilarity matrix were tested by running a PERMANOVA [79] via adonis2 and a PERMDISP [80] via betadisper (vegan v2.6 [81]). Differentially abundant taxa and/or functions were identified with ANCOM-BC 2 [82]. Enrichment/depletion in COG categories and/pathways within differentially abundant functions was tested with a hypergeometric test via phyper (stats v4.2.2). Throughout the analyses, a Holm– Bonferroni adjusted *p*-value < 0.05 was considered statistically significant, except for the ANCOM-BC 2 analyses, where a Holm–Bonferroni adjusted *p*-value < 0.001 was used.

### Data availability

All the FASTQ sequence data and associated metadata have been deposited in NCBI’s Sequence Read Archive under accession no. PRJNA995784. All the analyses conducted in this manuscript and additional data tables are available at https://github.com/CUMoellerLab/Real_etal_2023_MUPKO.

## Acknowledgments

Funding was provided by the National Institutes of Health (NIH) grant R35 GM138284-01 (A.H.M.) a pilot grant award under NIH Animal Models for the Social Dimensions of Health and Aging Research Network R24 AG065172 (M.J.S). NIH had no role in study design, data collection and interpretation, or the decision to submit the work for publication. The authors declare no competing interests.

We acknowledge Cornell’s Stem Cell and Transgenic Core Facility for generating the *Mup*-knockout mice and Cornell’s Biotechnology Resource Center for the Ilumina NextSeq 2000 sequencing. We also thank Tess Reichard for her help with the urine collections and the SDS-PAGE, and Dr. Weiwei Yang and Dr. Jon Sanders for their assistance with the mouse fecal DNA extractions, DNA library preparation, and submission of the DNA libraries for sequencing.

M.V.F.R., Conceptualization, Methodology, Investigation, Formal analysis, Visualization, Writing – original draft, Writing – review & editing; M.S.C., Methodology, Investigation; M.J.S., Conceptualization, Methodology, Investigation, Writing – review & editing, Funding acquisition; A.H.M., Conceptualization, Methodology, Investigation, Formal analysis, Visualization, Writing – original draft, Writing – review & editing, Funding acquisition.

## Supplemental Results

To investigate the effect of sampling bias on the detection of significant effects of *Mup* deletion on the microbiota in males but not in females, we subsampled our WT males by alternatively removing one male from the analyzes and testing for differences in microbiota beta and alpha diversity and differential abundance at both taxonomic and functional (COG) profiles. PERMANOVA analyses testing for differences in species composition in the gut microbiota of WT and KO males revealed that significance was maintained in 3/4 cases based on Jaccard dissimilarities, and in 1/4 cases using Bray-Curtis dissimilarities (Table S7). In terms of COG-function composition, significance was maintained in 2/4 subsampling analyses based on Bray-Curtis dissimilarities but not maintained for analyses based on Jaccard dissimilarities (Table S7). Significant differences in microbial family Shannon diversity were observed for 1/4 cases (Table S8), but not for alpha diversity measures based on COG functions (Table S8). Importantly, subsampling had no qualitative effect on the detection of differentially abundant taxa or functions in males. The significant effects of *Mup* deletion in males observed in these subsampling analyses—particularly the male-specific differential taxon and COG-function results, which were completely robust to subsampling—lend further support to our conclusions that *Mup* deletion more strongly affected the microbiota of male mice than female mice.

## Supplemental Figures

**Figure S1.**
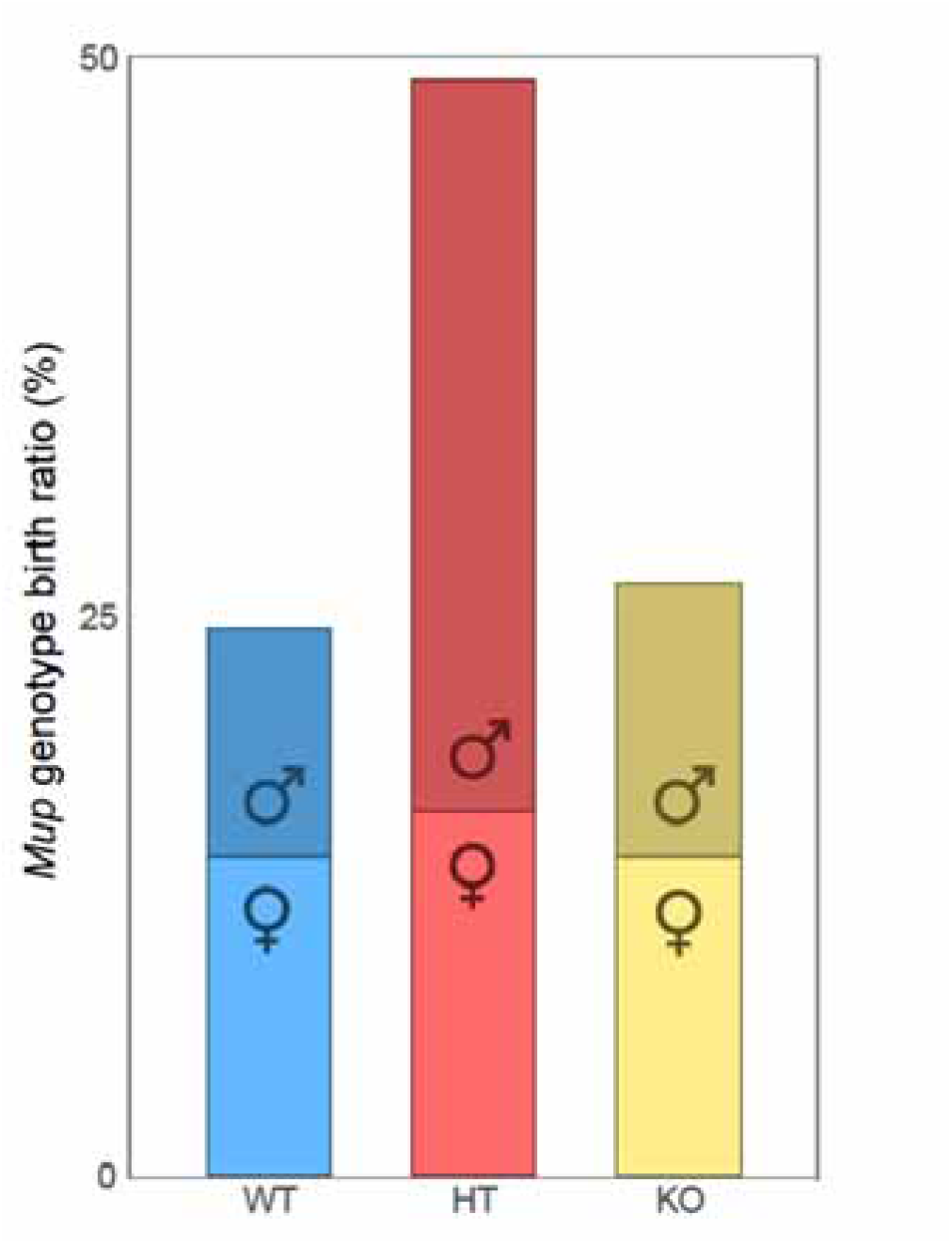
*Mup* genotype birth ratio. Bar plot shows the total percentage of male (top bar) and female (bottom bar) wildtype (blue), heterozygote (red), and knockout (yellow) pups born in the six study litters.

**Figure S2.**
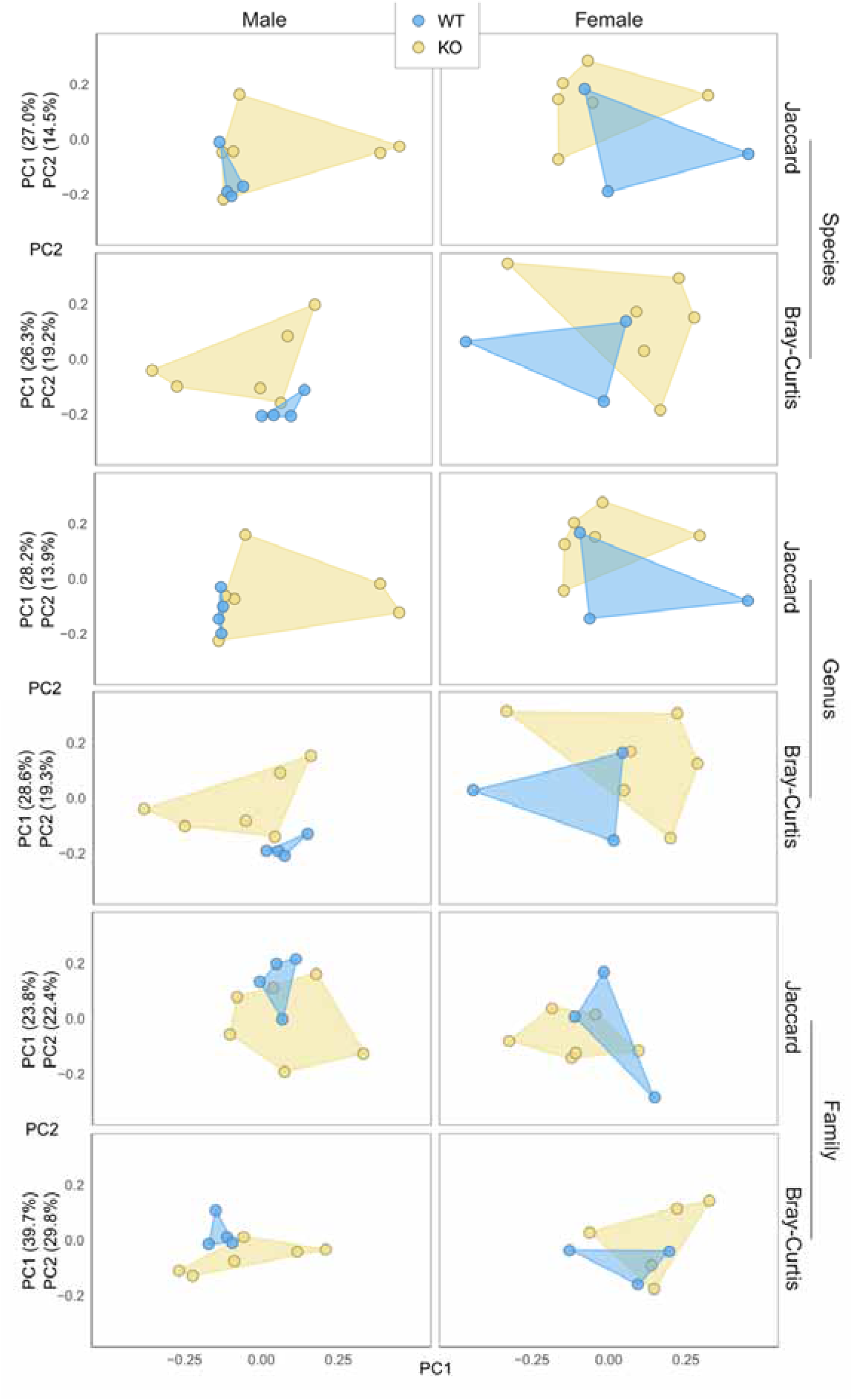
*Mup* deletion significantly changes the gut microbial taxonomic composition of mature males. Principal coordinates analyses (PCoA) show the ordinated Jaccard and Bray-Curtis dissimilarities in Species, Genus, and Family composition in the fecal metagenome of sexually mature mice. The ordinated points are faceted by sex and colored by genotype, showing the taxonomic profiles of WT (blue) and KO (yellow) male (left column) and female (right column) mice. The percentage of variation in the taxonomic dissimilarity matrix explained by the first two PCo axes is enclosed in parenthesis.

**Figure S3.**
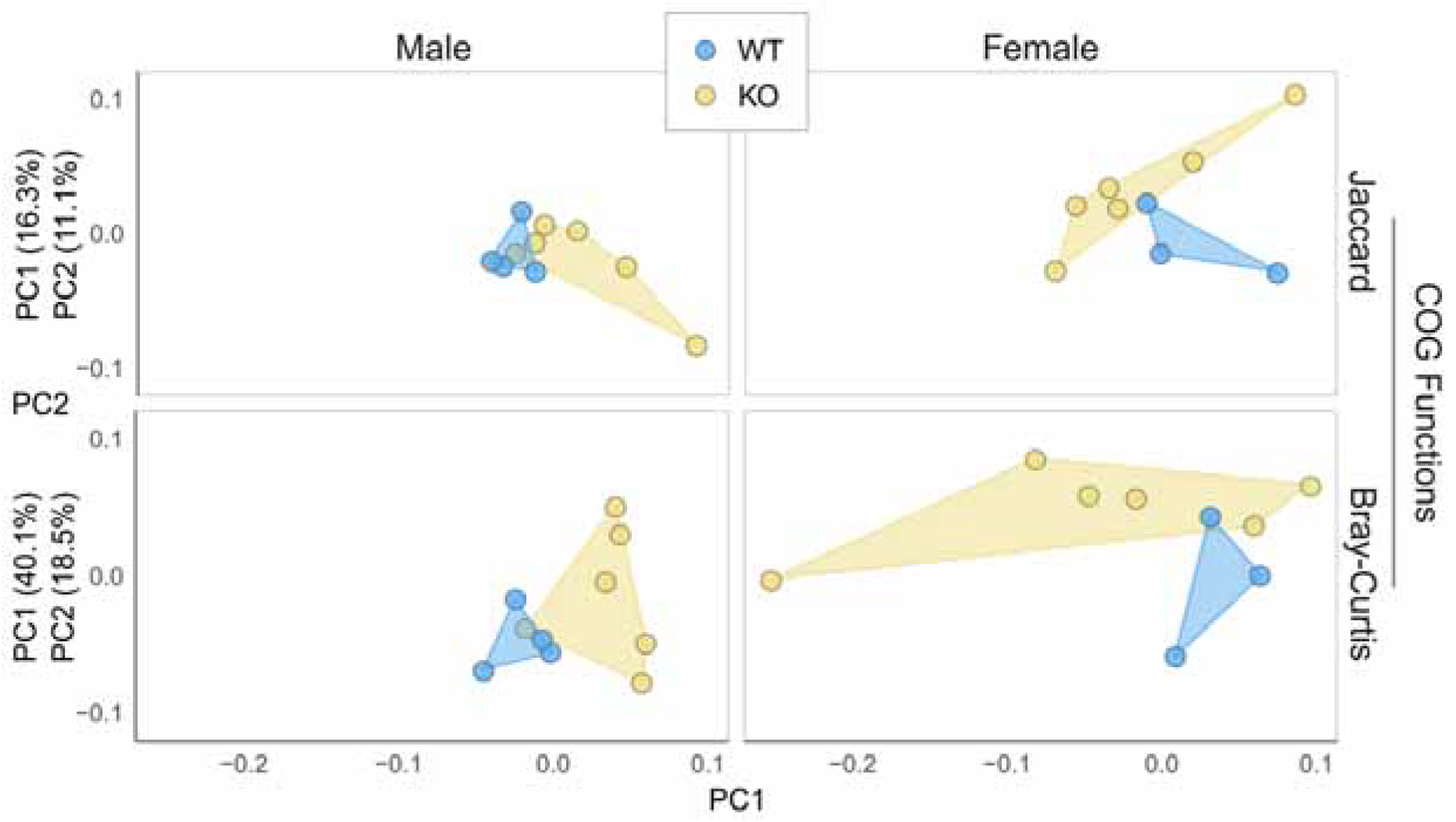
*Mup* deletion significantly changes the gut microbial functional composition of mature males. Principal coordinates analyses (PCoA) show the ordinated Jaccard and Bray-Curtis dissimilarities in COG Functions composition in the fecal metagenome of sexually mature mice. The ordinated points are faceted by sex and colored by genotype, showing the functional profiles of WT (blue) and KO (yellow) male (left column) and female (right column) mice. The percentage of variation in the taxonomic dissimilarity matrix explained by the first two PCo axes is enclosed in parenthesis.

**Figure S4.**
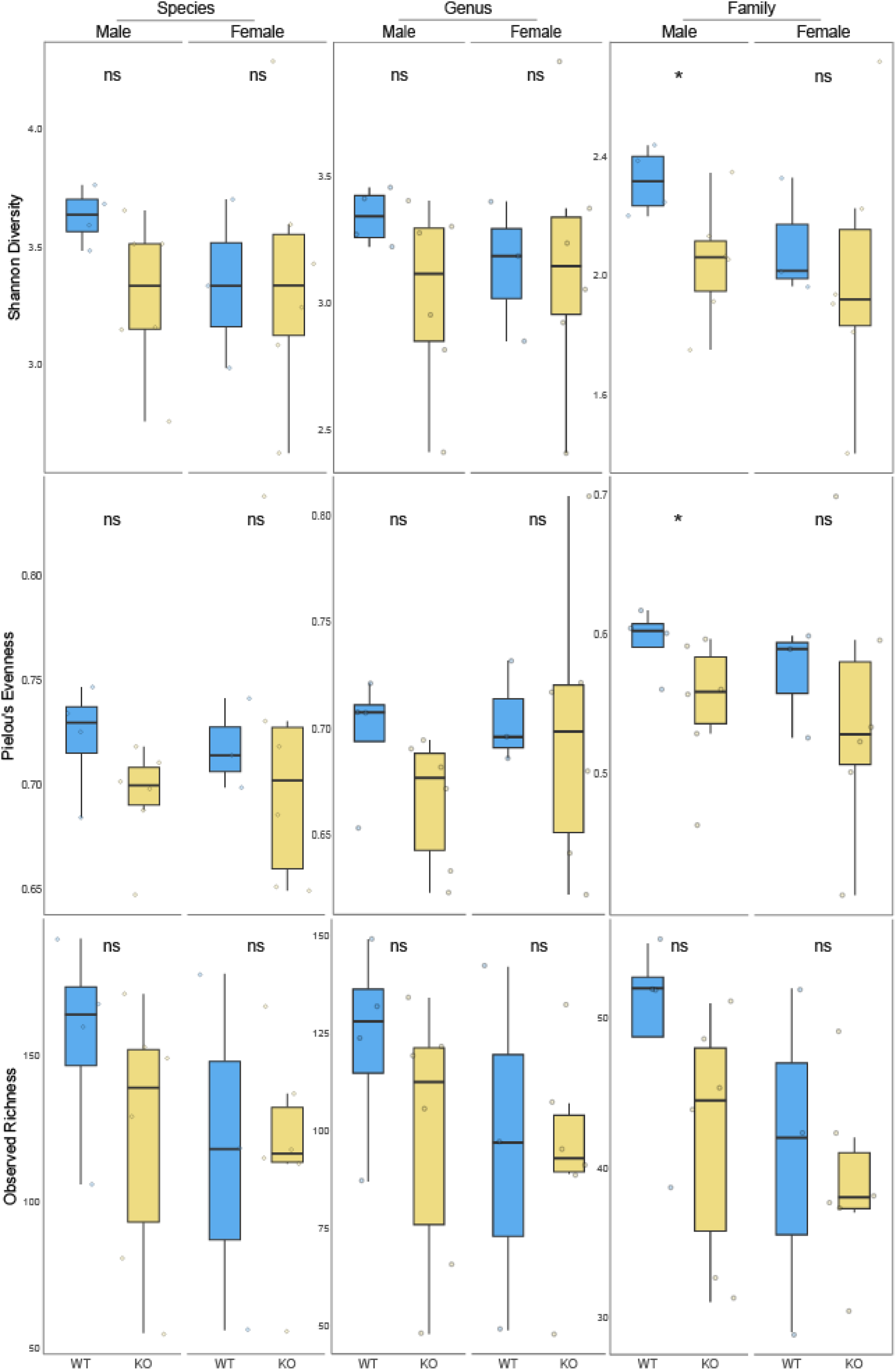
*Mup* deletion significantly reduces microbial family diversity in mature males. Box plots show the Shannon Diversity (top row), Pielou’s Evenness (middle row), and Observed Richness (bottom row) in microbial species (left column), genus (middle column), and families (right column) in the gut microbiota of mature males (left sub-column) and females (right sub-column). The linear mixed-effects model analyses (center top) indicate whether there is a significant difference in diversity in the gut microbiota of the WT (blue) and KO (yellow) mice (* = *p*-value < 0.05; ns = *p*-value > 0.05).

**Figure S5.**
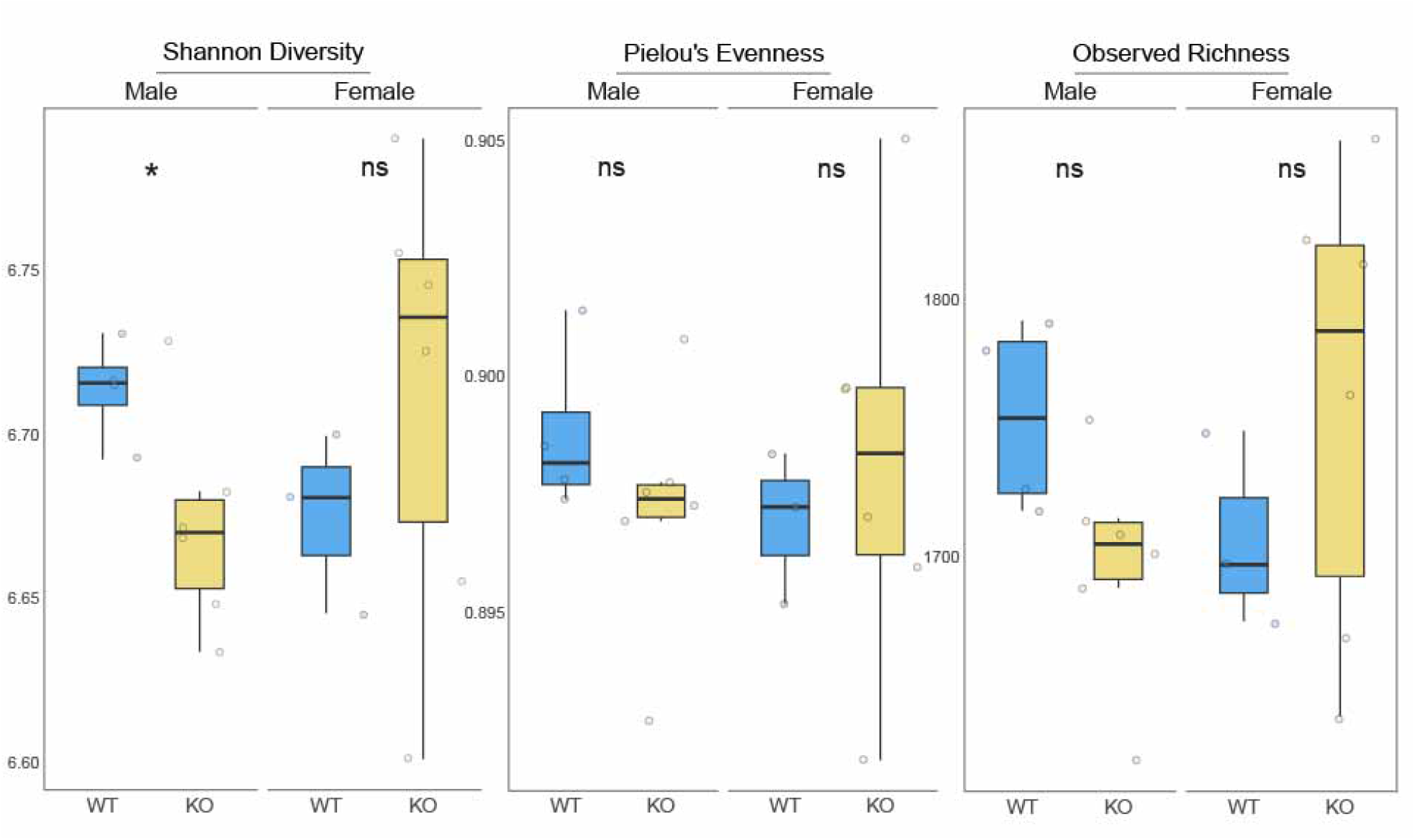
*Mup* deletion significantly reduces microbial functional diversity in mature males. Box plots show the Shannon Diversity (left column), Pielou’s Evenness (middle column), and Observed Richness (right column) in microbial COG Functions in the gut microbiota of mature males (left sub-column) and females (right sub-column). The linear mixed-effects model analyses (center top) indicate whether there is a significant difference in diversity in the gut microbiota of the WT (blue) and KO (yellow) mice (* = *p*-value < 0.05; ns = *p*-value > 0.05).

**Figure S6.**
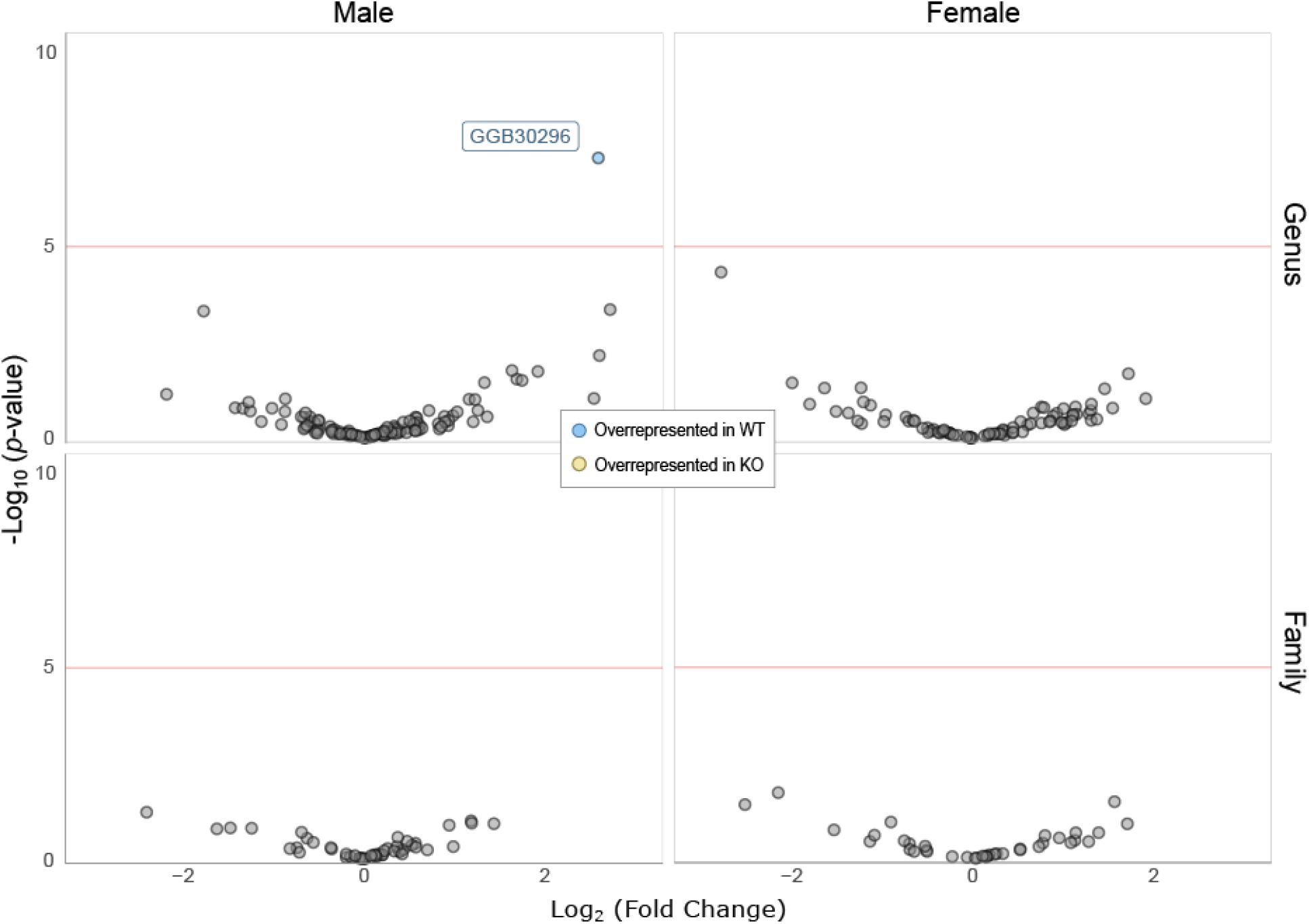
*Mup* deletion significantly shifts the abundance of various microbial taxa. Volcano plots show the log_2_-transformed fold change (LFC) in the abundance of genus (top row) and families (bottom row) in the gut microbiota of mature male (left column) and female (right column) mice. ANCOM-BC2 analyses identified taxa (points) that were significantly more abundant in WT (blue) or KO mice (yellow). Red lines mark the significance threshold (Holm–Bonferroni–adjusted *p*-value < 0.001). The y-axis indicates the -log_10_ transformation of the non-adjusted *p*-value.

## Supplemental Tables

**Table S1.**
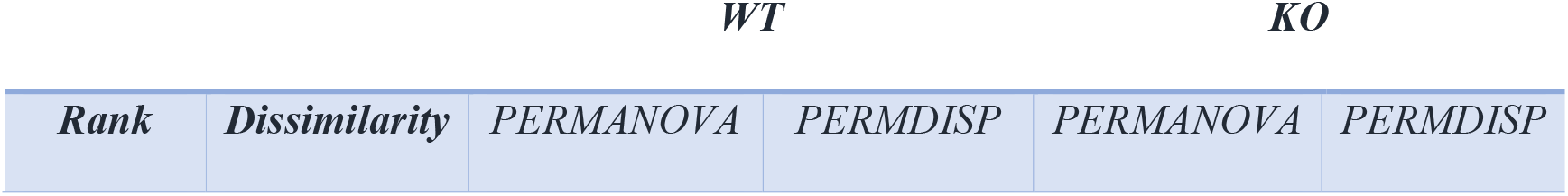
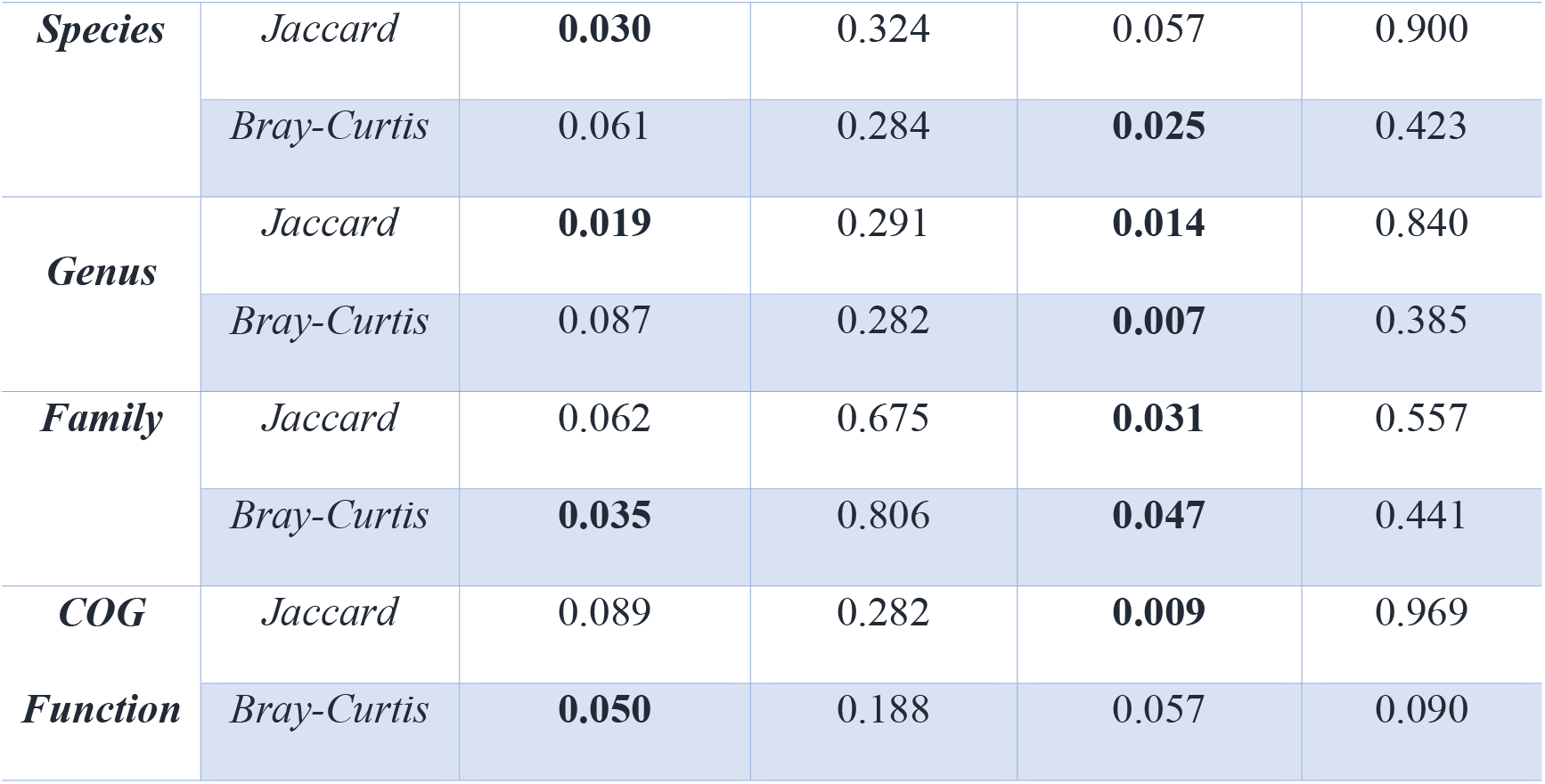
Effect of host sex on microbial taxonomic and functional composition (*p*-value).

**Table S2.**
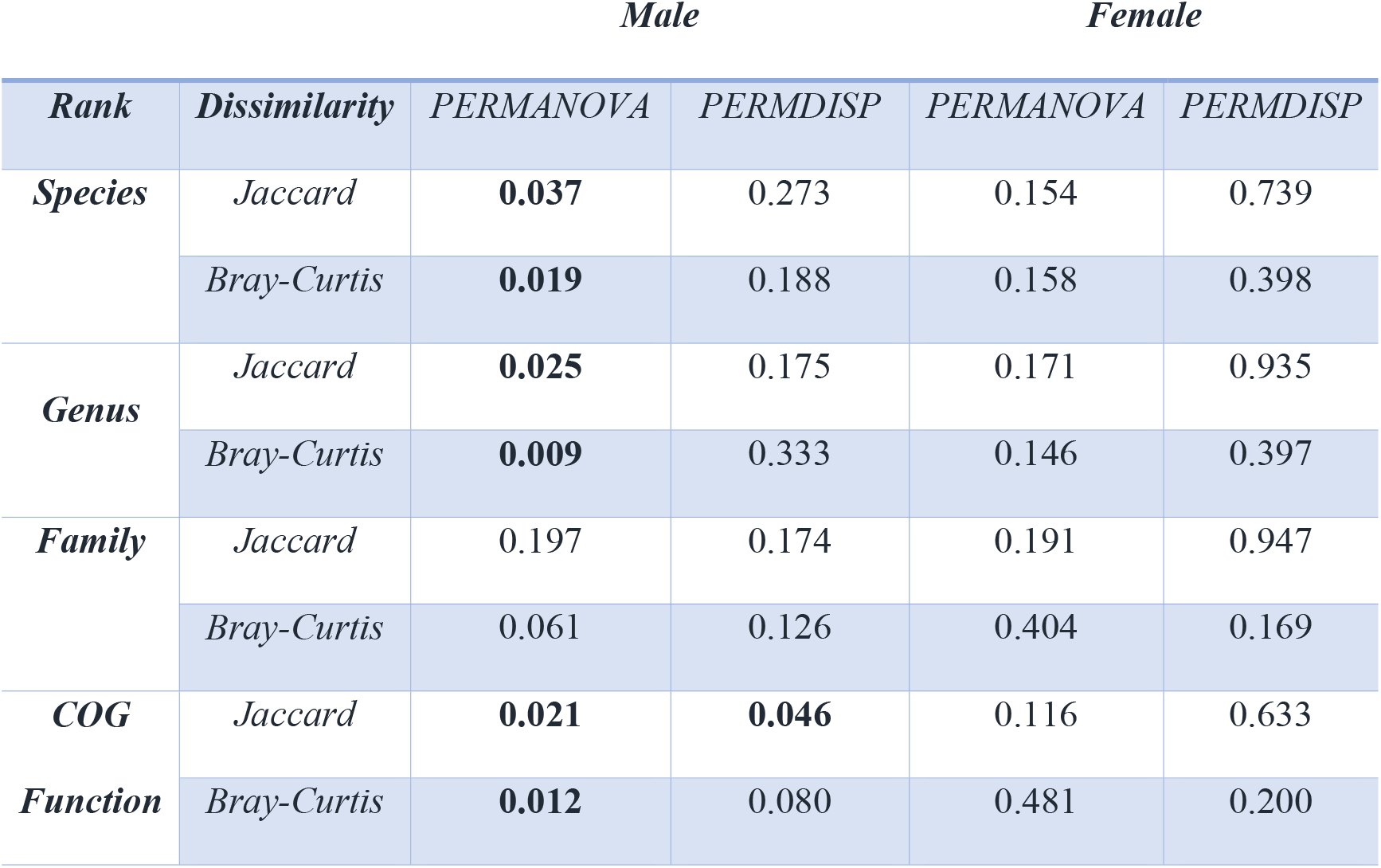
Effect of *Mup* genotype on microbial taxonomic and functional composition (*p*-value).

|  |  | *Male* |  | *Female* |  |
| --- | --- | --- | --- | --- | --- |
| *Rank* | *Dissimilarity* | *PERMANOVA* | *PERMDISP* | *PERMANOVA* | *PERMDISP* |
| **Species** | *Jaccard* | **0.037** | 0.273 | 0.154 | 0.739 |
|  | *Bray-Curtis* | **0.019** | 0.188 | 0.158 | 0.398 |
| **Genus** | *Jaccard* | **0.025** | 0.175 | 0.171 | 0.935 |
|  | *Bray-Curtis* | **0.009** | 0.333 | 0.146 | 0.397 |
| **Family** | *Jaccard* | 0.197 | 0.174 | 0.191 | 0.947 |
|  | *Bray-Curtis* | 0.061 | 0.126 | 0.404 | 0.169 |
| **COG** | *Jaccard* | **0.021** | **0.046** | 0.116 | 0.633 |
| **Function** | *Bray-Curtis* | **0.012** | 0.080 | 0.481 | 0.200 |

**Table S3.** Microbial taxonomic and functional profiles correspondence with Procrustes (*p*-value).

| *Rank* | *Dissimilarity* | *Male* | *Female* |
| --- | --- | --- | --- |
| ***Species + COG*** | *Jaccard* | **0.002** | 0.953 |
|  | *Bray-Curtis* | **0.001** | 0.230 |
| ***Genus + COG*** | *Jaccard* | **0.001** | 0.943 |
|  | *Bray-Curtis* | **0.001** | 0.221 |
| ***Family + COG*** | *Jaccard* | **0.001** | 0.866 |
|  | *Bray-Curtis* | **0.001** | 0.920 |

**Table S4.**
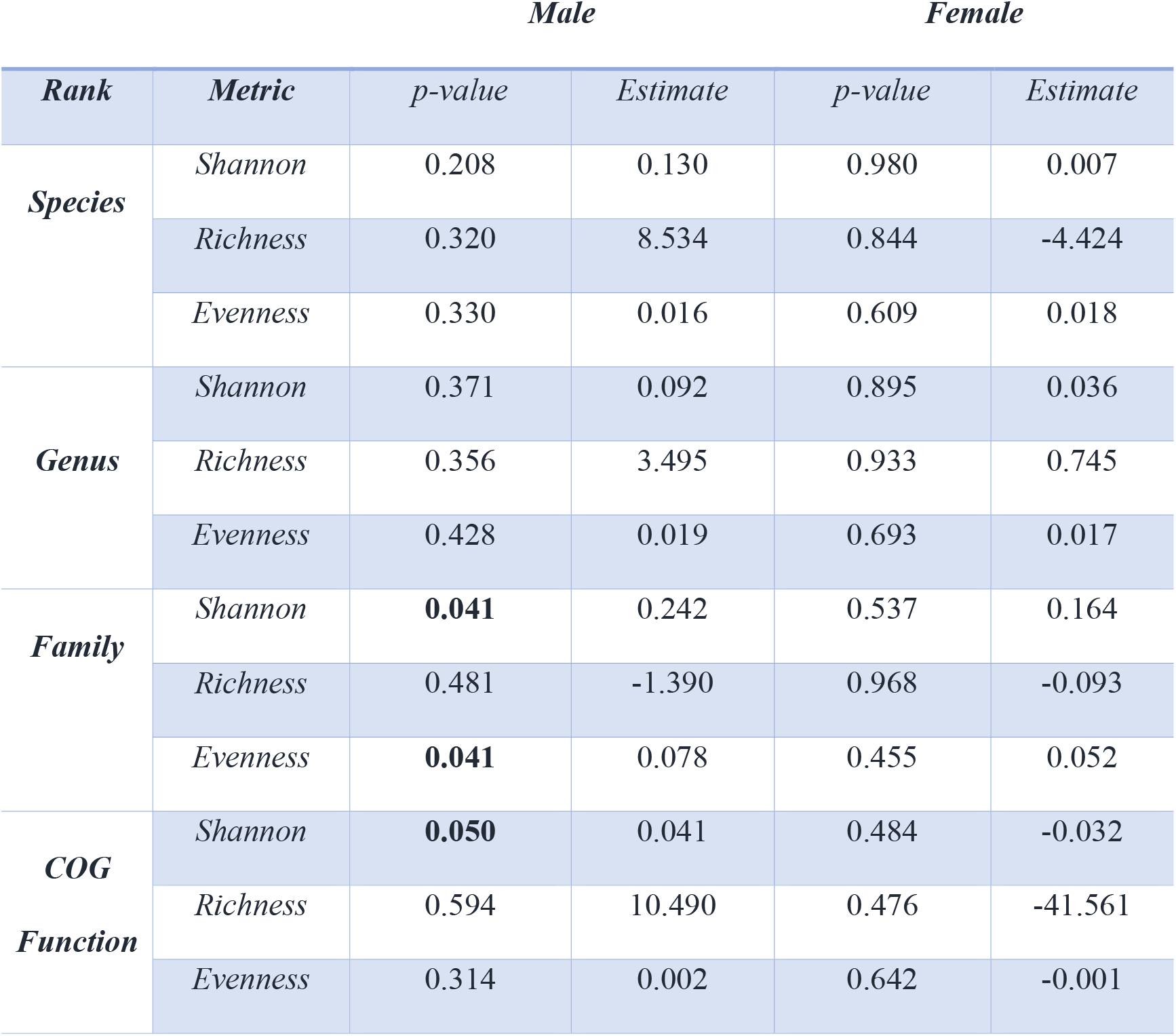
Effect of *Mup* genotype on microbial taxonomic and functional diversity (*p*-value).

**Table S5.**
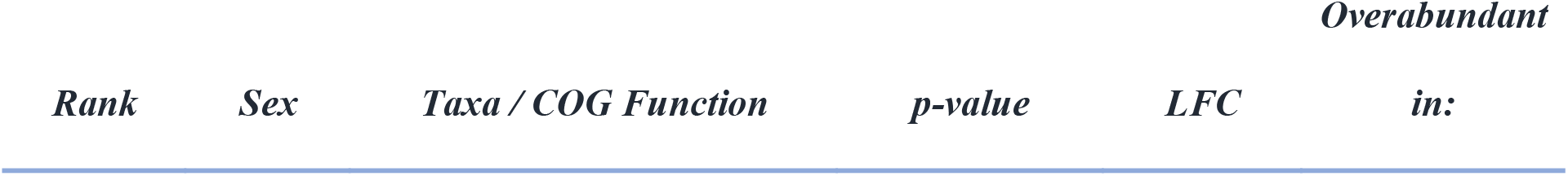
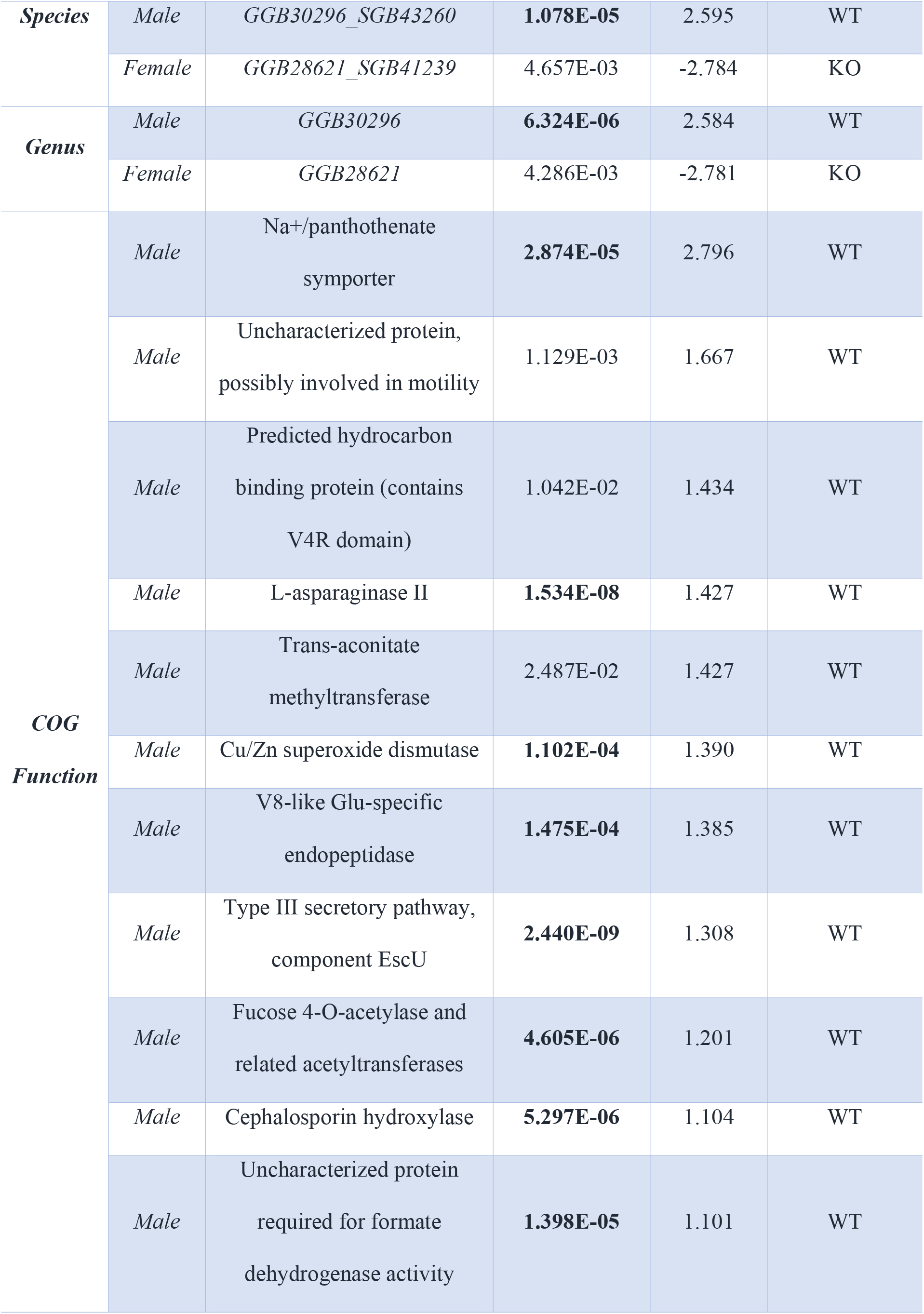
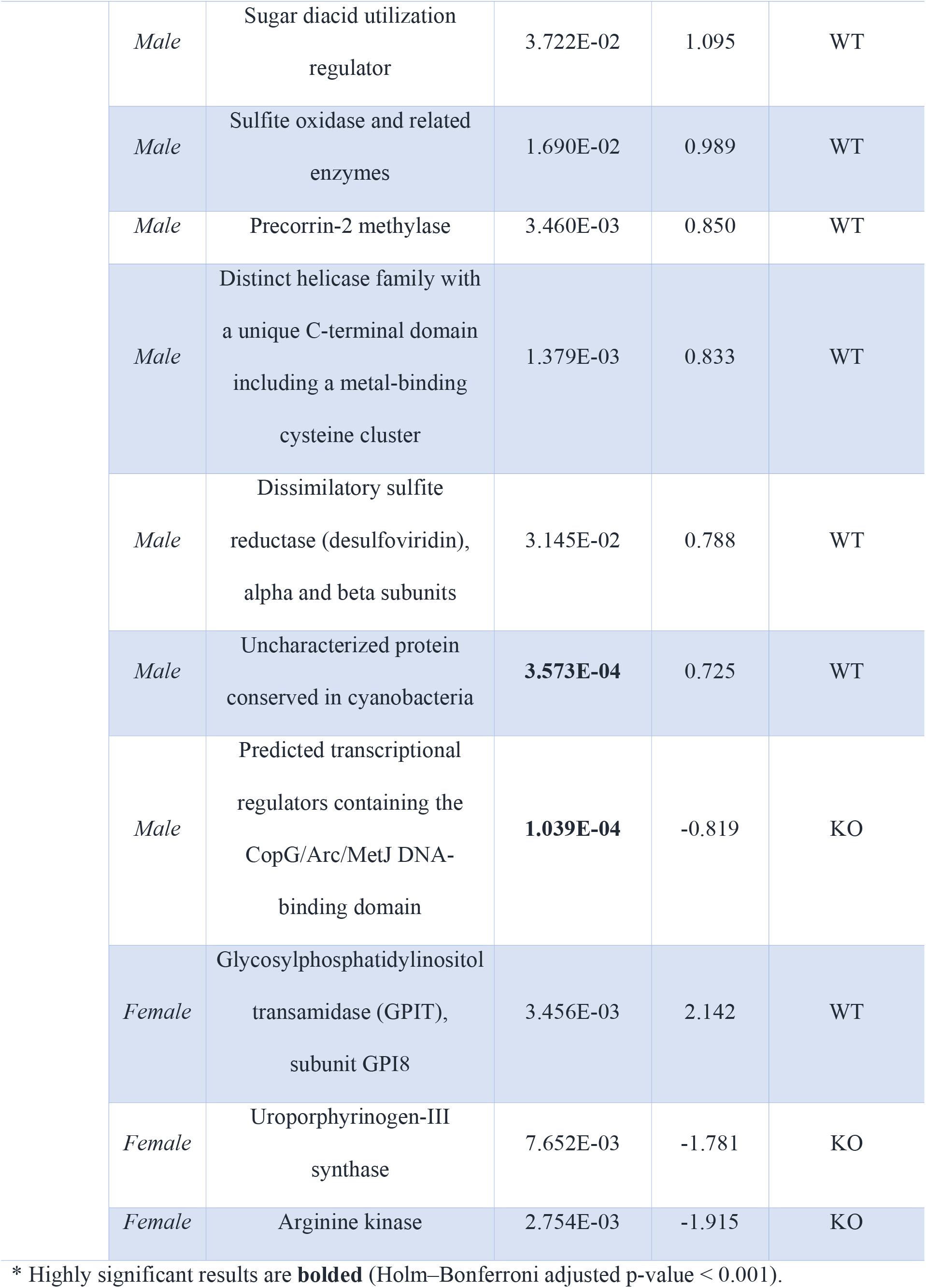
Microbial taxa and functions showing differential abundance in WT vs. KO genotypes*.

**Table S6.** Functional enrichment analyses with a hypergeometric test (*p*-value).

| *COG Category* | *Male* |  | *Female* |  |
| --- | --- | --- | --- | --- |
|  | *WT* | *KO* | *WT* | *KO* |
| *Cellular processes and Signaling* | 0.545 | 1.000 | 0.205 | 1.000 |
| *Information storage and Processing* | 1.000 | 0.170 | 1.000 | 1.000 |
| *Metabolism* | 0.425 | 1.000 | 1.000 | 0.189 |
| *Poorly characterized* | 0.251 | 1.000 | 1.000 | 1.000 |

**Table S7.** Power analysis of taxonomic and functional beta diversity results (*p*-value).

| *Rank* | *Dissimilarity* | *w/o 219* | *w/o 224* | *w/o 256* | *w/o 257* |
| --- | --- | --- | --- | --- | --- |
| *Species* | *Jaccard* | **0.014** | **0.039** | 0.058 | **0.041** |
|  | *Bray-Curtis* | 0.119 | 0.069 | 0.153 | **0.046** |
| *Genus* | *Jaccard* | 0.052 | 0.549 | 0.118 | 0.097 |
|  | *Bray-Curtis* | **0.017** | 0.097 | 0.077 | **0.029** |
| *Family* | *Jaccard* | 0.320 | 0.138 | 0.100 | 0.431 |
|  | *Bray-Curtis* | 0.084 | 0.126 | 0.053 | 0.175 |
| *COG* | *Jaccard* | 0.051 | 0.089 | 0.108 | 0.171 |
| *Function* | *Bray-Curtis* | **0.032** | **0.018** | 0.052 | 0.120 |

**Table S8.** Power analysis of taxonomic and functional alpha diversity results (*p*-value).

| *Rank* | *Metric* | *w/o 219* | *w/o 224* | *w/o 256* | *w/o 257* |
| --- | --- | --- | --- | --- | --- |
| *Species* | *Shannon* | 0.209 | 0.806 | 0.567 | 0.565 |
|  | *Richness* | 0.613 | 0.053 | 0.685 | 0.677 |
|  | *Evenness* | 0.259 | 0.102 | **0.024** | **0.034** |
| **Genus** | *Shannon* | 0.331 | 0.567 | 0.859 | 0.868 |
|  | *Richness* | 0.795 | **0.050** | 0.522 | 0.516 |
|  | *Evenness* | 0.252 | 0.138 | **0.025** | **0.027** |
| **Family** | *Shannon* | **0.034** | 0.184 | 0.077 | 0.117 |
|  | *Richness* | 0.185 | 0.084 | 0.629 | 0.494 |
|  | *Evenness* | 0.145 | 0.229 | 0.095 | 0.133 |
| **COG**<br>**Function** | *Shannon* | 0.087 | 0.057 | 0.102 | 0.152 |
|  | *Richness* | 0.373 | 0.559 | 0.558 | 0.546 |
|  | *Evenness* | 0.802 | 0.141 | 0.303 | 0.399 |

